# Evolutionary dynamics of maximal syllable complexity show no universal bias rooted in jaw biomechanics

**DOI:** 10.64898/2026.10.15.759367

**Authors:** Laura C. Dees, Shelece M. Easterday, Paul Widmer, Balthasar Bickel, Chundra A. Cathcart

## Abstract

Linguistic evolution has given rise to variation in the degree of syllable complexity that any given language permits. However, several models of language converge in predicting a universal bias towards the consonant-vowel (CV) structure (or a broader strong-weak asymmetry). This is closely tied to incremental speech processing, and in evolutionary terms has been linked to certain rhythmic orofacial displays in non-human primates. Yet, whether linguistic evolution really reflects such a universal tendency is an open question, one that we aim to address using phylogenetic comparative methods. Modeling change according to a continuous-time Markov process, we estimate transition rates for maximal onset and coda sizes (i.e. grammatical rules) across 5 language families. We demonstrate that a lineage-specific model outperforms a global model, and further explore the nature of lineage-specific long-term preferences and rates of change on multiple levels. We show that the former drive inter-family variation more than the latter. Generally, particulars turn out to impact syllable complexity more than any universal biomechanically or neurocognitively motivated biases do, implying that evolutionary continuity with our primate ancestors may have been overstated for this domain of language.

## 1 Introduction

Linguistic evolution has given rise to a wide spectrum of syllable complexity across the approximately 7000 languages spoken today, with complexity here meaning the maximal structure a given language allows. Maximal syllable structure in spoken languages ranges from simple (only consonant-vowel syllables such as *ba*) to highly complex. Languages of the latter type allow three, four or more consonants (depending on their nature) consecutively in the syllable beginning and/or end, often with additional combinatorial constraints. Despite this cross-linguistic diversity, evidence from various approaches to language, including neurobiology, converges on predicting a bias towards the simplest consonant-vowel (CV) structure, and towards “strong” syllable beginnings (onsets) and “weak” syllable ends (codas) more broadly. In addition, a growing body of comparative literature finds support for a degree of evolutionary continuity – the roots of syllabic speech run deep within the primate lineage. Here we ask whether the supposed bias is present as a factor in linguistic evolution. We use phylogenetic comparative methods to model the evolution of syllable complexity in grammar across human languages. We first briefly describe the different lines of evidence that directly or indirectly point towards a CV bias.

In literature on phonological theory and typology, the CV structure has received qualifications such as “basic” or “fundamental” [1], “optimal” [2], “preferred” and “least marked” [3, 4]. These labels are heavily inspired by the worldwide distribution of syllable structure: CV is universally present in the sense that it is the only structure that every language has in its inventory, thus lying at the basis of an implicational hierarchy [5, 3]. If a language allows codas at all, the onset tends to harbor a larger portion of the consonant inventory, larger clusters and/or fewer combinatorial constraints [4, 6, 1]. Apart from syllable inventories on the level of grammar, CV is also claimed to predominate in usage within and across languages [1, 7, 8] and onset clusters tend to be more frequent than coda clusters in languages that allow both [6]. Linguistic processes that seem to sustain the onset-coda asymmetry are phonological consonant strengthening domain-initially and weakening domain-finally [9] and phonetic consonant lengthening word-initially [10].

In parallel to the findings in linguistic theory, change and global distributions, several models of speech processing converge in predicting a CV bias (or the looser onset-coda asymmetry). From the perspective of speech perception, various findings point in the same direction. For example, in contrast to syllables with codas, CV offers clearest transitional cues to a consonant’s identity [11]. This benefits auditory recoverability of segments [1]. A similar advantage is offered by the maximized modulations of high (vowels) and low (consonants) intensity in CV structures [1], a view that matches older ideas on sonority sequencing in speech [12, 13]. The prominence of syllable and word onsets relative to codas has since long played a central role in literature on the incremental nature of speech processing and chunking [14]. Onsets have been found to be easier for listeners to identify than codas [15] and are more informative than following sounds [16, 14].

Production-related arguments for a CV bias have received perhaps even more attention, with more far-reaching evolutionary implications. Studies on articulatory gestures have found a discrepancy in the timing of onset versus coda consonants, with onset-vowel combinations being more stable and synchronous than vowel-coda combinations [17, 18], thus relating a CV bias to motor control. Pushing a production-related origin further back, the highly influential Frame/Content theory of MacNeilage [19] proposes that the ubiquity of CV is the direct outcome of phylogenetically old mechanical and temporal constraints on oscillatory movements of the jaw, with CV being the natural outcome of open-close alternations. These jaw oscillations are taken to have evolved from ingestion-related rhythms found in mammals that were co-opted as the “frame” for syllables. An apparent intermediate stage can be observed in communicative orofacial visual and/or audio signals (most prominently lip-smacking) in a range of non-human primates. Complementary evidence from language ontogeny suggests that babbling in human infants is subject to the same biomechanical constraints, supporting this theory [20, 21].

This evolutionary scenario is consistent with work on the neurobiological foundations of speech rhythm and its origins within the primate lineage. In the context of growing evidence for evolutionary continuity not only in vocal anatomy [22] but also in neuroanatomy and neurophysiology [23, 24], ample support has emerged for a potential homology between primate facial displays featuring mandibular oscillations and the universal open-close alternations of articulatory gestures in human language. Both occur within the theta frequency range and show similarities in the coordination of the vocal tract articulators, the involved neural architecture and their ontogenetic development [25, 26, 27, 28, 29, 30, 31, 32, 33]. The coupling of phonatory and articulatory (mandibular) oscillations in marmoset vocalizations makes for a particularly strong connection [33]. In turn, human-oriented neurobiological work stresses how the highly regular temporal structure of speech at the syllable level is exploited in perception and production systems, and is a consequence of the underlying neural foundations and coordinated movements of the jaw, lips, tongue and velum [34]. In sum, the open-close cyclicity that characterizes human speech rhythm is widely held to be subject to biomechanical constraints on our anatomy that have evolutionary origins deep within the primate lineage.

Together, these findings suggest that the syllable plays a central role in processing [34]. This in itself is reason enough to expect strong and universal selectional pressures towards an optimal CV structure in language. But the idea of such a pressure is in apparent conflict with the variation in syllable complexity observed across the globe [5, 35]. This calls into question how the current distribution in languages could evolve. Indeed, recent findings cast doubt on the universality of a CV preference: work on specific languages [36] or areas [37, 38] implies that perhaps the ubiquity of an onset-coda asymmetry has been overstated. Additionally, higher complexity can apparently remain diachronically stable within families or areas for a long time and does not seem to get lost easily [39].

A more basic limitation, however, is that synchronic distributions can offer only limited insight into evolutionary trajectories. Since apparent synchronic preferences do not necessarily reflect evolutionary biases, it remains an open question whether diachronic change favors the predicted CV bias (or the weaker onset-coda asymmetry). We aim at resolving this question by phylogenetically modeling the evolutionary dynamics of syllable complexity according to a continuous-time Markov process. We consider language-level maximally allowed syllable structure to gain insight into the domain of grammar. With this approach we infer transition rates and test through model comparison whether the evolution of syllable complexity can best be captured as a universal profile or by family-level dynamics. Findings indicate overwhelming support for the latter. We further explore family-specific trends in the domain of long-term preferences, gain and loss rates and system-level speed of change to characterize how different parameters variably contribute to diversification.

## 2 Results

### 2.1 Model comparison

We compare the goodness of fit of two phylogenetic models, a global model in which changes in onset/coda complexity occur according to a single process of evolution shared across all families in our sample of data, and a family-specific one in which this process varies across families in a restricted fashion via a hierarchical model. Bayes Factors provide decisive support (well above the consensual threshold of 100 [40]) for the family-specific model for both onsets (BF ≈ 8.55 × 10^8^) and codas (BF = 23708). This indicates that there is virtually no support for the idea that change in syllable complexity is characterized by the same process of diachronic change across families.

### 2.2 Long-term preferences for syllable complexity levels

We further inspect parameters of the family-specific model in order to investigate lineage-level long-term preferences and dynamics of change. We quantify family-level long-term preferences via the stationary probability of each family-level continuous-time Markov (CTM) process, which represents the limiting behavior of the process, i.e., the proportion of time that system is expected to be in different states of the process as time approaches infinity. These preferences can thus be regarded as independent of time and, as opposed to sole frequency distributions, are potentially informative of cognitive, communicative or functional pressures in linguistic evolution [41, 42]. The results are visualized in Fig. 1. In the domain of the onset, we find no cross-family preference for maximal onsets of size 1, nor for any other particular state: only three families (Austronesian, Pama-Nyungan and Uto-Aztecan) show high stationary probabilities for this state. Sino-Tibetan on the long term tends towards biconsonantal onsets and Indo-European towards triconsonantal ones, with probabilities being slightly more distributed in these two families. A cross-family bias can likewise be ruled out for codas. Crucially, none of the families demonstrate a long-term preference for maximal codas of size 0, i.e. absence of codas is clearly dispreferred in all families. Austronesian, Sino-Tibetan and Uto-Aztecan show a long-term preference for coda size 1 that contrasts sharply with other attested states. Pama-Nyungan tends towards 1 as well but has broadly distributed stationary probabilities for state 2. Moderate degrees of complexity are preferred in Indo-European judging from the relatively high probabilities of coda sizes 2 and 3.

**Figure 1:**
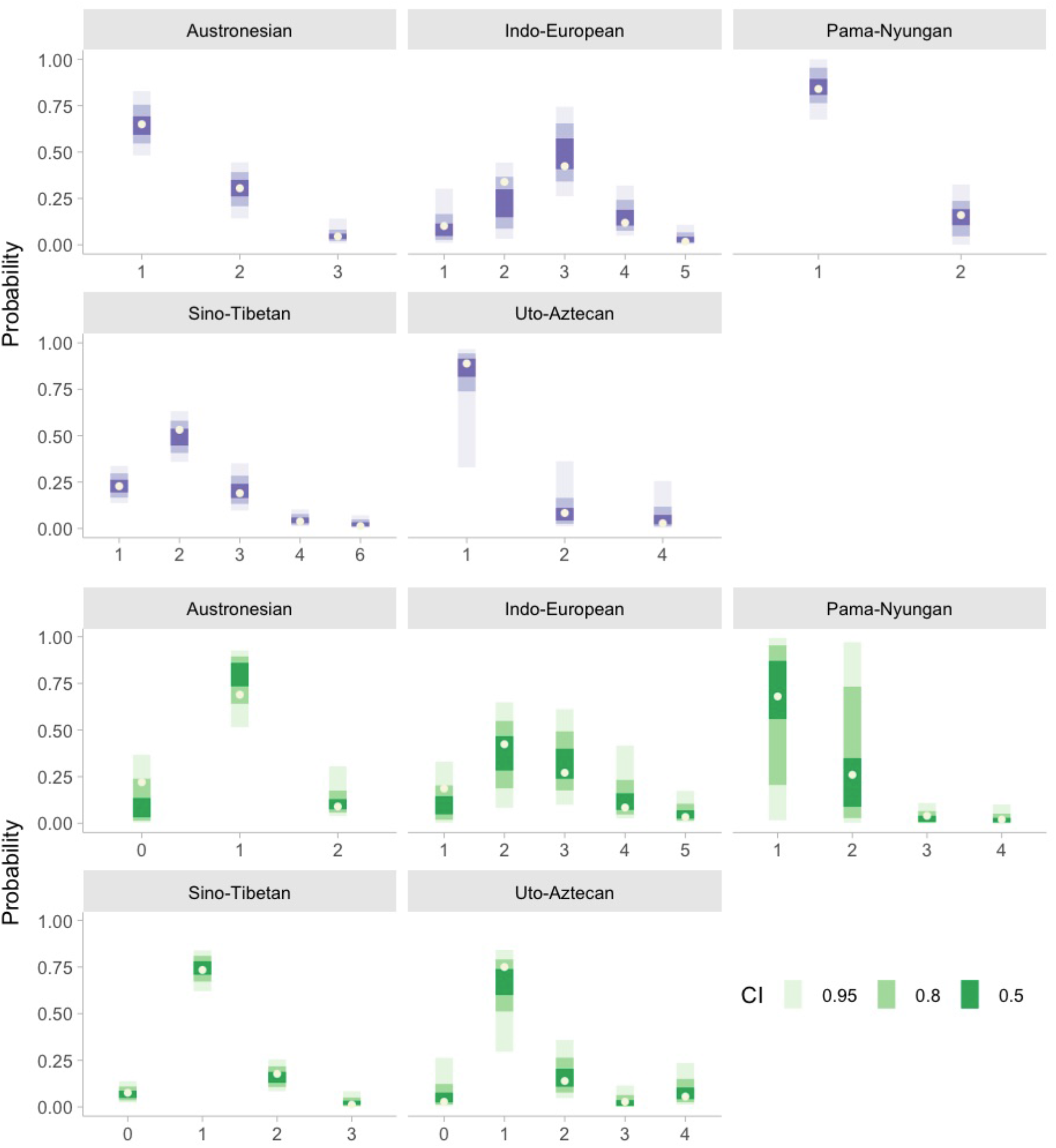
Family-level stationary probabilities represented by credible intervals for each of the ascertained maximal onset (upper half, in purple) and coda (lower half, in green) sizes. Observed probabilities are displayed in the form of beige dots within the intervals. For both the onset and coda, families show variation as to their preferred states, i.e. complexity levels with highest stationary probabilities. Predictions of the CV bias are not borne out across the five families.

In order to quantify the extent to which families vary in their stationary probabilities, even when they may prefer the same states on the long term, we calculated the pairwise differences between families for each onset/coda complexity level (see Supplementary Materials, Figs. 1-2). Naturally only states that occur in at least two families could be included. In the coda only 6 out of 32 family pairs demonstrate a decisive difference, all of which include Indo-European. In the onset we find 16/26 such pairs. These do not concentrate around any particular families. The picture that emerges from the stationary probabilities and the pairwise differences is that families show variation in their stationary probabilities for onset and coda sizes, though some families coincide in which states they prefer. In the case of coda states (that are attested in multiple families), families also show considerable overlap in the extent to which they prefer them. In the onset we observe substantial cross-family variability in stationary probabilities.

Comparing states with high stationary probabilities across onset and coda within each family through visual inspection, we observe that the predicted asymmetry in complexity is subtle – if present at all. Preferred onsets and codas are of the same size (1) in Austronesian, Pama-Nyungan and Uto-Aztecan. Only in Sino-Tibetan and Indo-European are the preferred onset clusters 1 consonant larger than the preferred codas (2 and 3 respectively for onset sizes).

### 2.3 Entry vs. exit rates

We next explore the pathways that lead to the long-term distributions, probing the possibility that families vary in the ways they attain and lose complexity levels. To this end we compare the rates at which onset and coda complexity levels are gained and lost across families (Fig. 2). For onset/coda states with high stationary probabilities, we expect high entry and low exit rates, and vice versa for states low stationary probabilities. Predictions seem to be only partly borne out in both the onset and coda.

**Figure 2:**
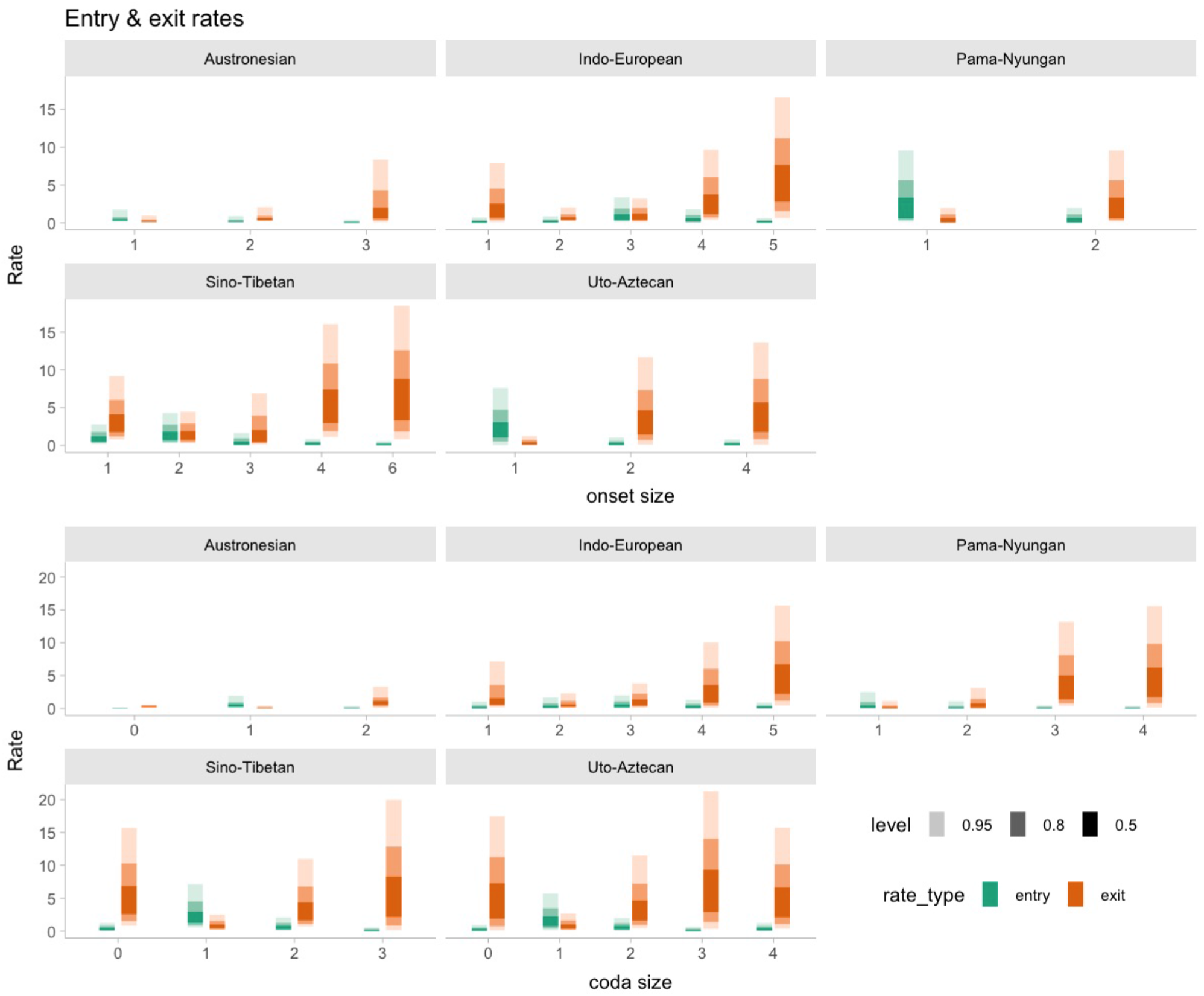
Family-level entry (green) and exit (red) rates represented by credible intervals for each of the ascertained maximal onset (upper half) and coda (lower half) sizes.

We base further judgment about between-family variation on posterior pairwise differences for each complexity level, for both entry and exit rates (see Supplementary Materials, Figs. 3-6). In the onset we find 6/26 family pairs with decisively different entry rates, and 3 such pairs in exit rates. In the coda, 4/32 family pairs demonstrate decisively different entry rates, and 5 for the exit rates. These data suggest that complexity levels are gained and lost at fairly similar rates across families, and therefore do not represent evidence that long-term preferences are heavily mediated by family-level discrepancies in entry and exit rates.

**Figure 3:**
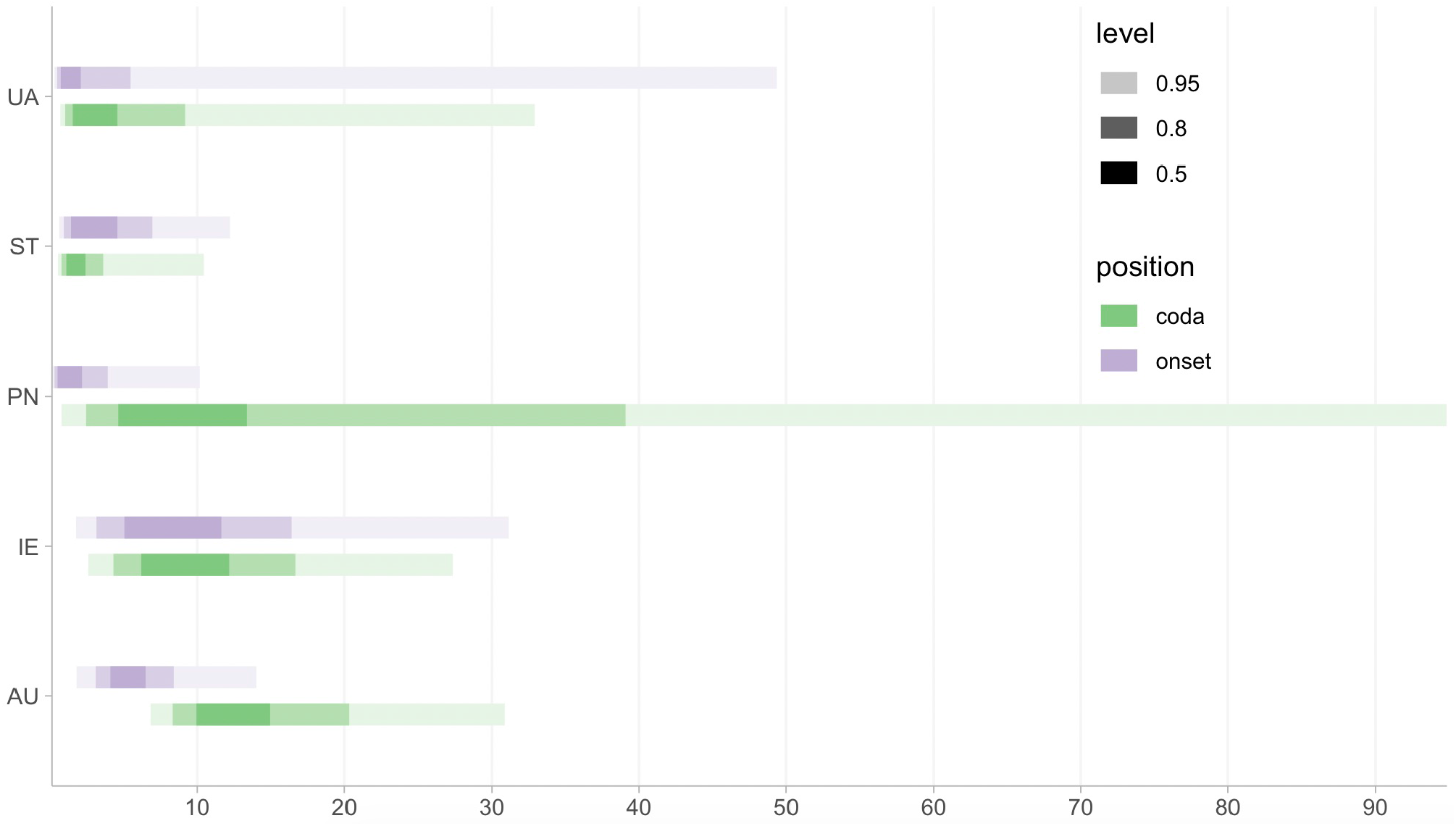
Family-level times to stationarity in millennia for maximal onsets (purple) and codas (green), represented by credible intervals.

Focusing on between-state differences we zoom in on the extent to which complexity levels vary in the way they are gained and lost *within* families. We examine pairwise posterior differences for both entry and exit rates (see Supplementary Materials, Figs. 7-10). These data amount to slightly more evidence for variation in the ways complexity evolves: out of 27 pairs of onset sizes, 10 are decisively different in the case of entry rates, and 7 in the exit rates. Pairs with robust differences in the coda add up to 8 out of 35 in entry as well as exit rates. Taken together, this does not provide strong evidence for substantial variation in the way complexity levels are entered and exited.

### 2.4 Speed of change

We compute the time to stationarity of each family-specific CTM process, i.e., the shortest time interval (in millennia) at which the transition probability distribution from all states converges on the stationary distribution of the CTM chain. As such, the distributions can be taken as a measure of overall speed of change in each specific family. Family-level posterior time-to-stationarity distributions are found in Fig. 3. Posterior pairwise differences reveal little support for substantial variation between families. We report 0 decisively different family pairs in the onset and 2 in the coda (see Supplementary Materials, Figs. 11-12). The latter indicate that Sino-Tibetan changes considerably faster than Austronesian and Indo-European.

Comparing onsets and codas visually, the inferred times to stationarity reported in Fig. 3 do not look particularly dissimilar within most families. To assess this formally we compared time-to-stationarity estimates in a meta-analysis of log0-transformed means where the posterior sample is treated as a Gaussian measurement error (Fig. 4). This suggests a weak tendency towards longer times to stationarity (i.e. slower overall rates of change) in codas compared to onsets, although uncertainty is substantial: the estimated difference includes zero in the 80% credible interval in four families and even in the 50% credible interval in one family. Put differently, we do not find evidence for the expected slower change in onsets.

**Figure 4:**
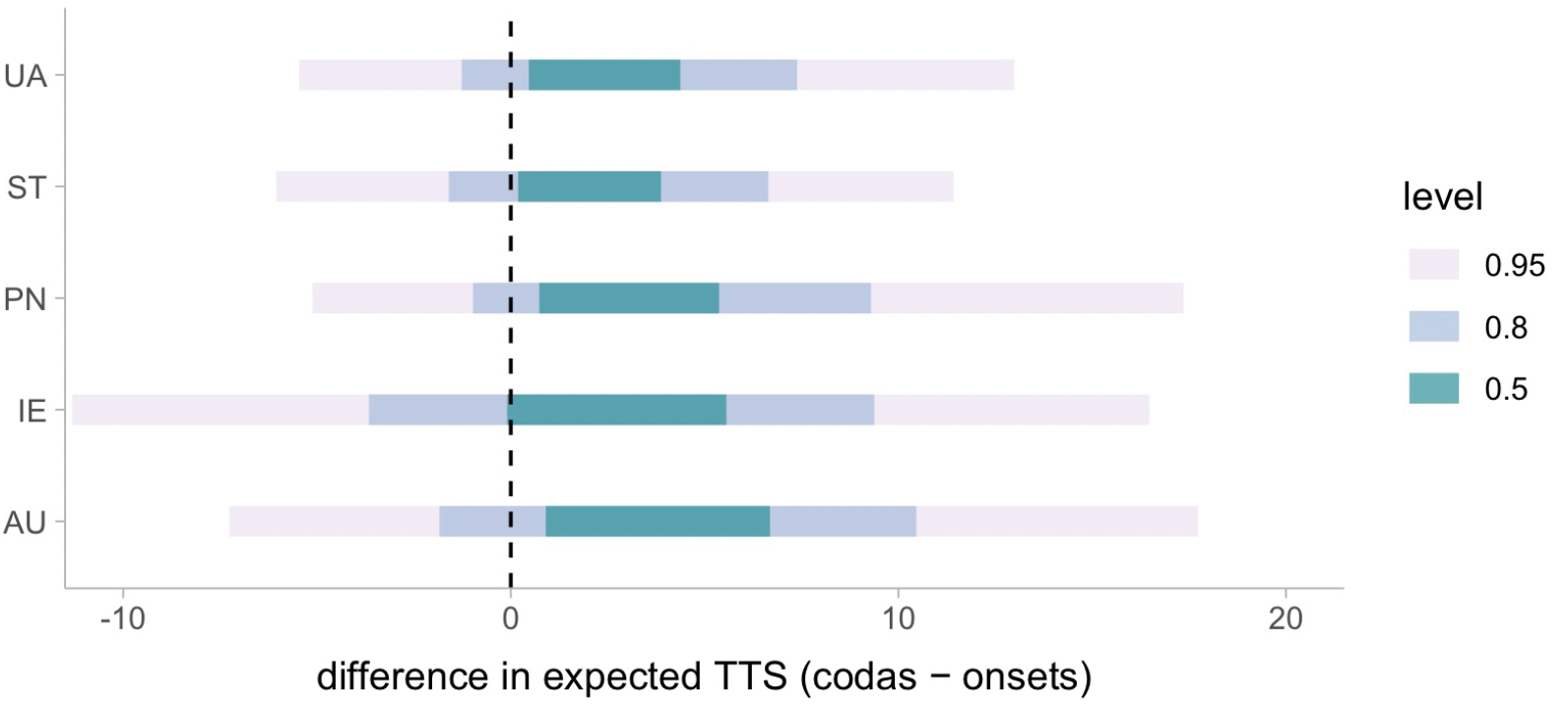
Posterior estimates of the difference in expected time to stationarity (TTS; codas − onsets) by family. Values greater than zero indicate longer times to stationarity in codas.

## 3 Discussion

Our model comparison shows that a lineage-specific model outperforms a global model in rate inference for both maximal onset and coda sizes. Notably, the lineage-specific model is favored despite (necessarily) having more parameters and Bayes Factors penalizing higher complexity. Further exploration of dynamics on the family level indicates that i) stationary probabilities speak against any cross-family long-term preferences for particular onset or coda sizes, with most pronounced cross-family variation in onsets, as quantified by pair-wise posterior differences; ii) entry/exit rates reveal little decisive variation, and more within families than between; iii) times to stationarity show no decisive family-level differences in onsets and only few in codas, and additionally no evidence for slower change in onsets compared to codas. Together, our findings indicate that maximal syllable complexity evolves according to overwhelmingly lineage-specific trends. These originate in differences in long-term preferences more than in variation in the tempo of change at the level of particular onset/coda sizes or the overall system.

Generally speaking, syllable complexity is modulated by a range of linguistic processes in the domains of phonology, prosody and morphosyntax. A comprehensive review of the immediate causes would lie outside the scope of this paper, and would in any case be limited by our insufficient understanding of both the mechanisms and the individual language histories. A brief look at the outliers is however warranted. In terms of speed, the faster change of codas in Sino-Tibetan compared to other families can be attributed at least in part to the development of tone: current evidence suggests that this happened many times independently and in slightly varying contexts, but with a key role for the loss of specific coda consonants (although onset consonants may be involved additionally or instead, depending on the language or subgroup [43, 44, 45]). As for the long-term preferences, Indo-European exhibits demonstrably higher coda complexity than other families. This is partly a matter of inheritance from the ancestral language, which already featured high complexity according to scholarly consensus. This preservation of complexity, but also loss with subsequent regeneration, can be linked to factors such as strong stress or accent systems with diachronic shifts that caused processes of vowel loss and subsequent cluster complexification, and an abundant ancestral system of suffixing inflectional morphology that is partly inherited into modern-day languages, all of which depends on the language or subgroup. The strong family variation in onset stationary probabilities and the consistent, though indecisive, evidence for faster speed of change in onsets than codas is surprising: mechanisms of sound change, with reductive processes often targeting non-initial positions, and the incremental nature of speech processing would predict more change-resistant onsets, therefore slower change. With this in mind, our result suggests that onset-preserving forces may be counterbalanced, decreasing the difference in speed with codas. While our understanding of the relevant linguistic processes does not allow for a purely mechanistic explanation, our tentative suggestion is that, perhaps, precisely because onsets are more salient to speakers and listeners in speech processing, they may be more contingent on the social factors that affect the pathways of change. Although the role of the pressure for social differentiation in linguistic evolution is far from understood, it does not seem inconceivable that such pressures impact onsets more than codas, thus speeding up change.

Crucially, the predictions made by a global CV bias are not borne out: the lineage-specific model speaks against cross-family uniformity in transition rates in the first place. Moreover, the inferred long-term preferences do not match with a general bias towards monoconsonantal onsets (although they are preferred in three out of five families) or absent codas (with very low stationary probabilities for 0). A weaker version of the CV bias predicts an asymmetry between onsets and codas, with onsets tending to be more complex. We find no robust evidence for this pattern: onsets are marginally more complex than codas in only two families and equally complex in the others. Importantly, these results apply to maximally allowed syllable structure, i.e. grammar, since the effects of a biomechanically or neurocognitively motivated bias would arguably reside there. If a CV bias operates primarily at the level of language use instead, it may still be a factor in the evolution of usage frequencies without leaving a trace in grammatical constraints.

The absence of a CV bias in our data can be interpreted in a broader evolutionary perspective. While the evolutionary roots of human motor action involved in syllable production may be shared with the rhythmic orofacial displays found in other primates, humans also developed a rich species-level sound repertoire, continuously evolving language-specific sound systems and extensive combinatorial capacities to recombine sounds into meaning-bearing units. Against this background, our results suggest that biomechanical or neurocognitive constraints on syllable complexity do not strongly limit current linguistic diversity. Factors that may have ultimately played a greater role in spurring diversity are of a sociocultural nature [23] – with a human urge for social indexing – and increased neural control over the vocal tract allowing for greater vocal flexibility. This aligns with recent calls [34, 25] to reconsider the sole importance of the jaw, and to place greater emphasis on how the coordinated movements of all vocal tract articulators shape speech rhythm processing. More work is needed to understand how the neurobiological underpinnings of speech rhythm map onto linguistic outcomes. More broadly, future work is needed to clarify how these sociocultural and anatomical factors interacted in the emergence of human phonological abilities and in the evolutionary dynamics underlying linguistic diversity today.

## 4 Materials and Methods

### 4.1 Syllable complexity data

The data consist of numerical values representing the maximal onset and coda structures, in number of consonants, for each individual language in our sample. We coded tautosyllabic onset (pre-vocalic) and coda (post-vocalic) clusters separately given that earlier findings in the literature predict different dynamics. Moreover, complexity levels in the onset and coda are coupled only to a limited degree and in a non-linear fashion [1, 39]. Data were manually coded in line with practices in Easterday [39] and extracted from reference grammars, either based on explicit statements, less explicit descriptions or, in rare cases, inferred from linguistic forms in the grammar at hand. We aimed to code maximal structures on the level of the phonological word domain [46]. If grammars used the more traditional notion of “word” instead, coding was based on that.

Hapaxes or low-frequency forms were considered relevant on the condition that they are invariant (i.e. there being no variant with a simpler syllable structure). In cases that exhibit variation, such as in (sociolinguistically determined) speech register, style and tempo, the more conservative variant was considered relevant. Loanwords were excluded if identifiable. No distinction was made between purely phonological clusters and clusters arising from morphological processes. Location in the word domain was also considered irrelevant. That is, our data do not distinguish between word-internal vs. word-edge clusters (though maximal syllable patterns correspond to the latter in most languages in the sample).

### 4.2 Phylogenetic data

The study spans 5 typologically and geographically diverse language families: Austronesian (represented in the sample by 177 languages), Indo-European (59), Pama-Nyungan (50), Sino-Tibetan (or “Trans-Himalayan”, 79) and Uto-Aztecan (36), rendering a total of 401 languages. These are almost exclusively contemporary varieties. Phylogenetic analyses were conducted on previously published posterior tree samples [47, 48, 49, 50, 51].

### 4.3 Methods

We employ phylogenetic comparative methods [52, 53, 54, 55] to characterize the dynamics of change between different degrees of onset and coda complexity. Evolution of coda and onset complexity are modeled separately. We model changes in onset/coda complexity over phylogenetic trees according to a continuous-time Markov (CTM) process, under which frequencies of change between states of a character — in our case, levels of complexity represented by onset/coda size — are characterized by non-negative transition rates. Our model allows direct transitions between non-adjacent as well as adjacent levels of complexity, as under some circumstances, sound changes such as vowel deletion can result in non-incremental changes in syllable complexity [39].

We compare the performance of two models: a global model in which transition rates are held constant across families and a family-specific model in which rates vary across families according to a hierarchical model. In the global model, change is modeled across all families according to a single CTM process characterized by transitions between all states attested across all families; in the local model, change is modeled in individual families according to family-specific CTM processes involving only transitions between states ascertained within those families. In each model, rates ***ρ*** ∼ Gamma(*α, α*), where the hyperparameter *α* ∼ Exponential(1). We compute family-level likelihoods under rate parameters and family-level phylogenies using the pruning algorithm [56]. Posterior distributions of rates are inferred using RStan [57]. In order to inject phylogenetic uncertainty into this procedure, we carry out rate inference for 25 trees from each family’s phylogenetic tree sample, aggregating posterior samples together. We ran models for 2000 iterations over 4 chains, discarding the first half of samples for each chain, resulting in 100000 posterior samples for each model (global vs. family-specific × onset vs. coda). Data were processed using version 2.1-1 of the R package phytools [58]. Data and code used in analyses can be found in the repository (*link available on publication*).

For both onsets and codas, we compare the fit of the global vs. family-specific models using Bayes Factors, a model comparison technique that quantifies the relative support for one model over another via a ratio of marginal likelihoods. We estimate model marginal likelihoods using version 1.1.2 of the R package bridgesampling [59]. Bayes Factors of greater than 100 are taken to represent decisive evidence in favor of the model represented by the numerator of the ratio.

For the family-specific models, we zoom in on different aspects of language evolution in line with recent work that aims to tease apart speed of change from long-term preferences [60, 42]. We calculate stationary probabilities of posterior CTM rate matrices using the R function svd. We estimate the time to stationarity of each family-specific posterior CTM rate matrix by finding the time interval (starting at zero and iteratively incrementing it with values of .01) at which the Kullback-Leibler divergence from the top row of the transition matrix to the stationary distribution drops below .01. To compare time to stationarity between onset and coda, we fit a hierarchical measurement-error model to log-transformed family-level mean times to stationarity. Position (onset vs. coda) was included as a predictor with varying intercepts and slopes by family, and family-level standard deviations were incorporated as measurement error. Models were fit using brms with Gaussian likelihoods and weakly informative priors. Posterior predictions were back-transformed to the original time scale, and differences in expected time to stationarity between codas and onsets were estimated by family.

For each posterior sample of rates, we compute the exit rate for state *i* by summing over all transition rates from *i* to other states, i.e., ∑ _*j≠i*_ *ρ*_*i→j*_ [61]. We compute the entry rate to a given state *i* following a formula derived by Gerhard Jäger *apud* [62], (∑ _*j ≠i*_ *π*_*j*_*ρ*_*j→i*_)*/*(∑ _*j≠i*_ *π*_*j*_), where ***π*** represents the stationary distribution of the CTM process characterized by rates ***ρ***. This results in posterior distributions of state-level exit and entry rates for each family.

When assessing the strength of differences between posterior parameters, we take there to be decisive evidence for a contrast if the 95% credible interval of paired differences across posterior samples excludes zero. For a one-tailed comparison, the 95% CI is bounded by the 5th to 100th percentiles; for a two-tailed comparison, the 95% CI is bounded by the 2.5th to 97.5th percentiles.

## Acknowledgements

The authors gratefully acknowledge the technical support provided by Carlo Y. Meloni as well as valuable feedback from audiences at SLE 2025 in Bordeaux and EVOLANG 2026 in Plovdiv. This research was funded by Swiss National Science Foundation grant no. 197389 and further supported by the NCCR Evolving Language (SNSF Agreement #51NF40 180888).

## Author contributions

Data curation and investigation were done by LD, SE and CC. LD, BB and CC designed the methodology and performed formal analyses. LD produced the visualizations and wrote the initial draft with detailed methodological input from CC. Funding was acquired by PW and BB. The project was administered by LD and PW. PW, BB and CC supervised the project. All authors contributed to conceptualization of the project and reviewing and editing the manuscript.

